# Determination of diagnostic cycle threshold (Ct) cut-offs for qPCR-based prevalence surveys of soil-transmitted helminth infections

**DOI:** 10.64898/2026.01.05.697616

**Authors:** Malathi Manuel, Nils Pilotte, Joseph W.S. Timothy, Sean R. Galagan, Gideon John Israel, Craig T Connors, Victor Omballa, Monica Pechanec Voss, Ushashi C Dadwal, Andrew M Gonzalez, Justine Ahlonsou, David Chaima, Zayina Zondervenni Manoharan, Doug Rains, Kristjana H. Ásbjörnsdóttir, Steven A Williams, Adrian J F Luty, Moudachirou Ibikounlé, Khumbo Kalua, Robin Bailey, Rachel L Pullan, Judd L Walson, Sitara SR Ajjampur

## Abstract

WHO guidelines for control of soil-transmitted helminths (STH) rely on coproscopic methods to assess population prevalence. In low-prevalence and light-intensity STH settings, quantitative PCR (qPCR) has higher sensitivity and specificity for detection. For qPCR to accurately identify transmissible infections of public health significance, it is essential to interpret the qPCR cycle threshold (Ct) results. As part of the DeWorm3 community-based cluster randomized trial on interrupting STH transmission, we conducted population-based surveys using high-throughput qPCR and aimed to establish appropriate Ct cut-offs to detect transmissible infections. Experimental approaches including egg and genome-equivalent spiking experiments were hindered by inefficient fecal DNA extraction despite optimization efforts. The Ct results for 29,980 samples (pre-intervention, cross-sectional surveys) revealed a bimodal distribution for two of the four species tested, *N. americanus* and *A. lumbricoides*. The first peak was assumed to represent transmissible infections, and the second peak to represent indeterminate or non-transmissible infections. Using a finite mixture model, we defined true qPCR positivity as any Ct result with a ≥5% chance of belonging to the first peak. This approach yielded Ct cut-offs of 34.4398 for *N. americanus* and 28.57587 for *A. lumbricoides*. For hookworms, sensitivity of qPCR was 96.7%, compared to 73.2% for Kato-Katz and moderate- to heavy-intensity infections (median Ct: 19.1, interquartile range [IQR]: 17.9-19.8) were differentiated from light-intensity infections and Kato-Katz negative samples (25.3, IQR: 22.5-27.9). Our findings demonstrate the feasibility and utility of evidence-based Ct cut-offs to identify transmissible STH infections in large scale surveys, and to categorize infection intensity as programmatically relevant.

**Author Summary:** Quantitative PCR (qPCR) has been used for the detection of soil-transmitted helminths but with limited emphasis on determining cycle threshold (Ct) cut-offs to accurately identify transmissible infections which are of public health significance. When qPCR is used to assess interventions, or, in the future, to potentially make programmatic decisions, it will be crucial to validate positivity criteria to avoid underestimation (false negatives) or overestimation (false positives) of results.

As part of the DeWorm3 trial, a community-based cluster randomized trial on interrupting transmission of STH, we developed and applied a validated STH qPCR to test 29,980 samples collected pre-intervention and explored experimental approaches to establish assay-specific Ct cut-offs. The Ct values from these samples showed a bimodal distribution for *N. americanus* and *A. lumbricoides*, suggesting the presence of two distinct groups. For this reason, a statistical approach with a finite mixture model was employed to determine Ct cut-offs that differentiated epidemiologically relevant, transmissible, egg-positive STH infections from those that are likely to represent detection of non-transmissible DNA or indeterminate results. While our data were applied in three different country settings, India, Benin and Malawi, it is important to note that no single Ct cut-off may be applicable across all epidemiological scenarios. Our findings demonstrate the feasibility of developing evidence-based Ct cut-offs with high sensitivity to accurately detect transmissible STH infections in large scale surveys.

## Introduction

Soil-transmitted helminths (STH, *Necator americanus, Ancylostoma duodenale, Ascaris lumbricoides,* and *Trichuris trichiura*) continue to be an important public health problem globally, with the most recent estimates suggesting that approximately 1.5 billion people, or 24% of the world’s population, remain at risk of infection [1]. The World Health Organization (WHO) strategy for STH focuses on controlling morbidity through targeted deworming of preschool-age (PSAC) and school-aged (SAC) children and women of reproductive age, by improving access to safe water, sanitation, and hygiene facilities, and by encouraging behavioural change [2]. In the last two decades, there has been a scale-up of targeted deworming activities with benzimidazoles in areas with moderate and high STH infection prevalence [3–5]. These control efforts have undoubtedly contributed to substantial reductions in the STH burden [6, 7] and will require large-scale impact assessments.

Coproscopic techniques, such as the Kato-Katz method, remain one of the commonly used tools in STH programmatic surveys to determine the appropriate frequency of targeted deworming [8]. However, these methods lack sensitivity in low-prevalence and light-infection intensity settings, making them poorly suited for impact assessment surveys after multiple rounds of deworming [9–11]. With improved diagnostic sensitivity, quantitative PCR (qPCR) represents a viable alternative to coproscopy [8, 11–12]. Crucially, however, modeling studies [13] have highlighted that specificity (rather than sensitivity) is of increasing importance as programs approach the endgame of morbidity elimination. In well-designed and adequately validated qPCR assays, where off-target amplification does not occur, a Ct (number of cycles) <40 is typically representative of a pathogen signal [14]. However, as qPCR-based assays detect parasite-derived target sequences regardless of viability, a positive signal may result from both viable eggs representing a transmissible infection as well as from non-viable DNA sources such as degraded, sloughed, or pass-through helminth material, thereby reducing specificity [15–17]. However, existing STH diagnostic studies have not established robust approaches to determine appropriate Ct cut-offs [18–21]. As qPCR methods are increasingly applied to STH programs, consideration must be given to how Ct results are interpreted to optimise both sensitivity and specificity.

As part of the DeWorm3 community-based cluster randomized trial, a multi-site qPCR assay protocol was validated [22], and utilizing a combination of experimental and statistical approaches, we have attempted to establish evidence-based qPCR assay Ct cut-offs for better estimation of epidemiologically relevant, transmissible STH infections. We also calculated the sensitivity and specificity of these assays compared to the Kato-Katz method.

## Methods

### Study setting and samples

As part of the DeWorm3 trial, a validated qPCR was carried out on 29,980 samples, results from which were used to determine appropriate qPCR Ct cut-offs for transmissible STH infections. Briefly, this trial was carried out in three sites - Benin, India, and Malawi - to test the feasibility of interruption of STH transmission by community-wide mass drug administration (MDA) [23]. Each study site was divided into 40 clusters, with at least 1,650 individuals per cluster and was randomized to receive six rounds of either bi-annual community-wide MDA with albendazole (20 clusters) or the standard-of-care national school-based deworming (SBD) program (20 clusters). In each site, the prevalence of STH was assessed by pre-intervention cross-sectional surveys (CS) (~20,000 stool samples collected per site) and post-intervention cross-sectional surveys (24 months post-MDA, ~40,000 stool samples collected per site). Additionally, stool samples were collected from a longitudinal monitoring cohort (LMC), pre-intervention for all sites (LMC1) and annually at selected sites (Benin and India), with samples comprised of 150 age stratified individuals from each cluster. All samples collected as part of the LMC1 were immediately tested by Kato-Katz in each site’s laboratory, the results of which have been reported previously [24]. After homogenization in 95% ethanol, all LMC and CS samples were stored at −80°C to ensure sample integrity and optimize DNA recovery [25].

### DNA extraction and qPCR

Semi-automated DNA extraction (with bead-beating) and qPCR were carried out using standardized protocols optimized and validated at testing laboratories as described [22]. Briefly, DNA was extracted with the MagMAX Microbiome Ultra Nucleic Acid Isolation Kit (Thermo Fisher Scientific), and qPCR performed using custom-designed lyophilized qPCR plates (Argonaut Manufacturing Services, Carlsbad, CA) containing a pair of triplex reactions (*Bacillus atrophaeus*, *N. americanus,* and *T. trichiura* and *B. atrophaeus, A. lumbricoides,* and *A. duodenale*) with STH primers targeting non-coding repetitive sequences [26, 27]. All samples were spiked with *B. atrophaeus* (ZeptoMetrix, Buffalo, NY) as an internal extraction and amplification control, and an *Ascaris suum*-spiked sample was included with each extraction batch as a positive control, along with a negative reagent only control. Testing of Benin and Malawi samples was carried out at Quantigen, Fishers, IN, United States, and samples from India were tested at Christian Medical College (CMC), Vellore, India (**S1 Fig**). To ensure harmonization of the qPCR protocol, robust quality assurance (QA) was implemented, whereby 10% of samples tested in each laboratory were retested in a different laboratory (10% of Benin and Malawi samples from Quantigen were sent to Institut de Recherche Clinique au Bénin (IRCB), Benin and CMC laboratories, respectively, and 10% of samples tested at CMC were sent to Quantigen). Each testing laboratory was periodically evaluated using a proficiency panel of four blinded samples from Smith College, Northampton, Massachusetts, United States, after every 4,600 study samples or 25 qPCR batches.

### Single egg spiking experiments

Spiking experiments were conducted at Quantigen using commercially available *Ascaris suum* eggs (Excelsior Sentinel Inc., Ithaca, NY), as this species is a near-identical genetic match to *A. lumbricoides* [28]. Additionally, *A. suum* eggs are hardy, presenting a suitable DNA extraction challenge. To prepare the samples, under a microscope, 1 µL of *A. suum* egg suspension (23 eggs/µL) was aliquoted onto a glass slide. Using a micropipette, single eggs were then added to 50 mg aliquots of naive stool sample. A total of 35 samples, each containing a single *A. suum* egg, were homogenized, resulting in ~800 µL of lysate per sample. In the first experiment, DNA extraction and qPCR was performed on 400 µL of stool lysate using the trial protocol [22] as well as by single-plex qPCR [27]. To ensure that the target was not missed due to sampling error, the remaining 400 µL of lysate also underwent extraction and was tested using single-plex *Ascaris* qPCR. In the second experiment, 800 µL of lysate was extracted in two 400 µL aliquots and the DNA was then pooled for analysis using the single-plex qPCR assay. In experiment three, increased bead beating was carried out prior to DNA extraction and qPCR testing (**S1 Table**).

### One genome equivalent spiking experiment

During embryonation, helminth ova contain a variable number of diploid cells; therefore, the number of genome copies may vary from egg to egg. To evaluate the impact of genome copy variability on Ct cut-off, we created and tested a panel of samples containing the minimum amount of DNA that could represent a single infectious egg (one diploid genome-equivalent) [26]. This panel consisted of 44 naive stool samples, each spiked with one diploid genome equivalent of DNA from three target species (*N. americanus*, *A. duodenale*, and *A. lumbricoides*). This experiment did not include *T. trichiura,* as the DNA material sourced was insufficient. DNA was extracted from each sample using the trial protocol and tested in triplicate using single-plex qPCR assays for each species [26, 27].

### Modeling of Ct cut-offs using finite mixture models

After pooling qPCR results from LMC1 and CS1 tested from all three sites, bimodal Ct distributions for both *N. americanus* and *A. lumbricoides* (**Fig 1**) were observed. This was not seen for *A. duodenale* and *T. trichiura* (which were of very low prevalence) nor for the *B. atrophaeus* internal control. Given the bimodal distributions observed for *N. americanus* and *A. lumbricoides* Ct values, we hypothesized that these peaks represented two underlying groups: transmissible, egg-positive infections and non-transmissible infections. Consequently, we fitted a bimodal finite mixture model (FMM) to the Ct density plots, enabling us to derive a Ct cut-off below which a result would be assumed to represent a transmissible infection. Finite-mixture models assume that an observed distribution is made of multiple, unobserved, subpopulations. These models leverage expectation-maximization algorithms to iteratively optimize the posterior likelihood of each sub-population and assign each observation a probability of belonging to each component, capturing both overlap and classification uncertainty.

**Fig 1.**
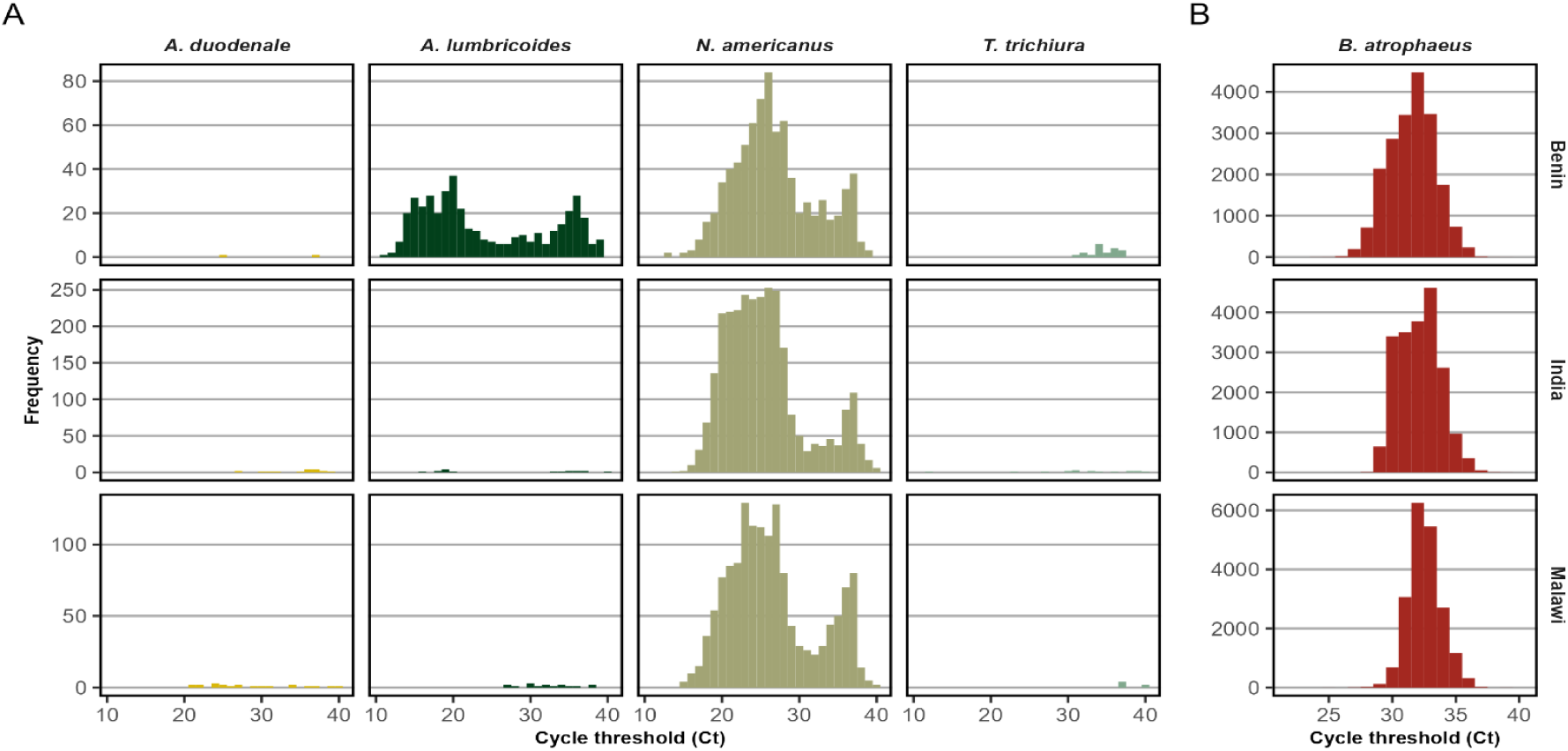
Distribution of Ct values for soil-transmitted helminth qPCR positive samples and *Bacillus atrophaeus* internal control qPCR positive samples from DeWorm3 Trial baseline surveys in Benin, India, and Malawi. (A) Ct values of STH positive samples (*A. duodenale, A. lumbricoides, N. americanus, T. trichiura*) when defining positivity as any amplification with Ct <40. (B) Ct values of *Bacillus atrophaeus* positive samples.

The qPCR results from LMC1 samples and a randomly selected subset of 20% of the CS1 samples from all three sites were used to inform the generation of Ct cut-offs. In the absence of strong prior expectations, we evaluated multiple candidate distributions (considering Gaussian, skew-normal, and t-distributions; and both equal and variable variance models) using the *mclust* package (version 6.0.0) in R Statistical Software (v4.2.2, R Foundation for Statistical Computing, Vienna, Austria) [29]. The best-fitting model was selected based on lower Bayesian information criterion (BIC). For *N. americanus*, FMM analysis was also conducted separately for each of the three study sites of varying prevalence to assess the consistency of the derived Ct cut-offs. Samples were subsequently classified as belonging to the transmissible egg-positive population if they had a ≥5% probability of belonging to the best-fitting egg-positive component; conversely, samples were classified as non-transmissible only if there was at least 95% probability of belonging to the best-fitting egg-negative, non-transmissible group. Samples with matching Kato-Katz results were then evaluated against these thresholds, allowing direct comparison of sensitivity and (mis)classification rates.

### Categorization of infection intensity by qPCR Ct

To classify infection intensity, the WHO classification was used (light-intensity infections were defined as <5,000 EPG for *A. lumbricoides* and <2,000 EPG for hookworm; moderate-intensity infections as 5,000 to 49,999 EPG for *A. lumbricoides* and 2,000 to 3,999 EPG for hookworm; and heavy-intensity infections as ≥ 50,000 EPG for *A. lumbricoides* and ≥ 4,000 EPG for hookworm [30]. For each infection intensity group, median Ct and interquartile ranges (IQR) were calculated, and the Kruskal-Wallis test was used to assess the differences in Ct distributions across the different infection intensity groups.

### Diagnostic accuracy of Kato-Katz and qPCR using Bayesian latent class analysis

Due to the absence of a gold standard diagnostic method, we estimated the sensitivity and specificity of Kato-Katz and qPCR using a Bayesian latent class analysis (BLCA). The BLCA was conducted using Markov Chain Monte Carlo (MCMC) methods via rjags package in R [31]. We modelled the observed diagnostic test outcomes across multiple populations under the assumption of conditional independence between Kato-Katz and qPCR results given true infection status. Non-informative uniform priors were used for all test parameter and population-specific prevalences. Model fit and parameter estimates were summarized by posterior means and 95% Bayesian Credible Intervals. Convergence diagnostics and effective sample sizes were used to assess model adequacy.

## Results

### Kato-Katz- and qPCR-based detection of STH in DeWorm3 baseline surveys

In Benin, India, and Malawi, a total of 6,089, 6,014, and 5,954 LMC1 samples, respectively, were tested by both Kato-Katz and qPCR. Additionally, 3,949, 3,979, and 3,939 CS1 samples were tested by qPCR alone from Benin, India, and Malawi, respectively (**Table 1**). When a qPCR Ct result of <40 was considered positive with no cut-off applied, the STH positivity in LMC1 samples was 11.7% by qPCR in Benin compared to 5.2% by Kato-Katz. In India, 27.1% of samples were positive for STH by qPCR compared to 17.0% by Kato-Katz, and in Malawi, the STH positivity was 14.9% by qPCR compared to 7.5% by Kato-Katz.

**Table 1.** Detection of soil-transmitted helminth infections by Kato-Katz, qPCR (without Ct cut-off), qPCR with genome equivalent derived Ct cut-off (GE+3SD) and qPCR with Finite Mixture Model (FMM) derived Ct cut-off.

| Site,<br>Activity | Samples tested | Any STH | Hookworm |  | Ascaris | Trichuris |
| --- | --- | --- | --- | --- | --- | --- |
|  |  |  | N. americanus | A. duodenale | A. lumbricoides | T. trichiura |
|  |  | n (%) | n (%) | n (%) | n (%) | n (%) |
| Benin |  |  |  |  |  |  |
| LMC1 | Kato-Katz (N = 6,089) | 319 (5.2) | 196 (3.2) |  | 124 (2.0) | 5 (0.1) |
|  | qPCR (N = 6,089) | 713 (11.7) | 483 (7.9) | 0 (0.0) | 246 (4.0) | 13 (0.2) |
|  | qPCR GE+3SD cut-off (N= 6,089) | 480 (7.9) | 303 (5.0) | 0 (0.0) | 179 (2.9) | - |
|  | qPCR (FMM cut-off) (N = 6,089) | 588 (9.7) | 427 (7.0) | - | 169 (2.8) | - |
| CS1 | qPCR (N = 3,949) | 477 (12.1) | 313 (7.9) | 2 (0.1) | 173 (4.4) | 6 (0.2) |
|  | qPCR GE+3SD cut-off (N= 3,949) | 328 (8.3) | 210 (5.3) | 2 (0.1) | 116 (2.9) | - |
|  | qPCR (FMM cut-off) (N = 3,949) | 376 (9.5) | 269 (6.8) | - | 108 (2.7) | - |
| India |  |  |  |  |  |  |
| LMC1 | Kato-Katz (N = 6,014) | 1,025 (17.0) | 1,004 (16.7) |  | 6 (0.1) | 17 (0.3) |
|  | qPCR (N = 6,014) | 1,632 (27.1) | 1,613 (26.8) | 12 (0.2) | 14 (0.2) | 11 (0.2) |
|  | qPCR GE+3SD cut-off (N= 6,014) | 1,245 (20.7) | 1,224 (20.4) | 12 (0.2) | 7 (0.1) | - |
|  | qPCR (FMM cut-off) (N = 6,014) | 1,472 (24.5) | 1,455 (24.2) | - | 7 (0.1) | - |
| CS1 | qPCR (N = 3,979) | 1,268 (31.9) | 1,261 (31.7) | 5 (0.1) | 3 (0.1) | 6 (0.2) |
|  | qPCR GE+3SD cut-off (N= 3,979) | 981 (24.7) | 973 (24.5) | 5 (0.1) | 1 (<0.1) | - |
|  | qPCR (FMM cut-off) (N = 3,979) | 1,130 (28.4) | 1,123 (28.2) | - | 1 (<0.1) | - |
| Malawi |  |  |  |  |  |  |
| LMC1 | Kato-Katz (N = 5,954) | 448 (7.5) | 431 (7.2) |  | 3 (0.1) | 15 (0.3) |
|  | qPCR (N = 5,954) | 885 (14.7) | 861 (14.5) | 14 (0.2) | 12 (0.2) | 6 (0.1) |
|  | qPCR GE+3SD cut-off (N= 5,954) | 643 (10.8) | 622 (10.5) | 14 (0.2) | 5 (0.1) | - |
|  | qPCR (FMM cut-off) (N = 5,954) | 776 (13.0) | 762 (12.8) | - | 2 (<0.1) | - |
| CS1 | qPCR (N = 3,939) | 587 (14.9) | 582 (14.8) | 7 (0.2) | 4 (1.0) | 0 (0.0) |
|  | qPCR GE+3SD cut-off (N= 3,939) | 355 (9.0) | 352 (8.9) | 7 (0.2) | 1 (<0.1) | - |
|  | qPCR (FMM cut-off) (N = 3,939) | 461 (11.7) | 458 (11.6) | - | 1 (<0.1) | - |

Footnote: Acronyms: LMC – Longitudinal Monitoring Cohort; CS – Cross-Sectional Survey; GE+3SD – Genome equivalent +3 standard deviations. For the genome equivalent-derived Ct cut-off (3SD), any STH prevalence was calculated by considering any amplification with Ct up to 40 as positive for *T. trichiura*, since this experiment was not conducted for *T. trichiura*. For the finite mixture model-derived Ct cut-off, overall STH prevalence was calculated by considering any amplification with Ct up to 40 as positive for *T. trichiura* and *A. duodenale*.

### Egg and genome spiking experiments to determine Ct cut-off

In the egg spiking experiment, only 31/70 (45.6%) of spiked samples tested positive for *A. suum/A. lumbricoides,* reflecting highly variable DNA extraction efficiency. This approach was not pursued further for Ct cut-off determination (**S1 Table**). In the genome equivalent spiking experiment, the mean Ct values for *N. americanus*, *A. duodenale*, and *A. lumbricoides* were 26.2, 33.1, and 28.8, respectively (**S2 Table**). Using these mean values, a three-standard deviation (3SD) Ct cut-off (i.e. capturing 99.9% of samples with one genome equivalent) was estimated as 27.8 for *N. americanus*, 39.7 for *A. duodenale*, and 30.6 for *A. lumbricoides*. Applying this 3SD cut-off, 72.7% of all 2,957 *N. americanus* qPCR positive samples (i.e. those with Ct<40 and a Kato-Katz test result) were classified as positive, including 93.2% of Kato-Katz positive samples (1411/1,514) and 51.1% of Kato-Katz negative samples (738/1,443). For *A. lumbricoides,* 70.2% of all samples with Ct<40 (with a Kato-Katz result) classified as positive with the 3SD cut-off (191/272), including 100% of Kato-Katz positive samples (109/109) and 50.3% of Kato-Katz negative samples (82/163). Applying this 3SD cut-off to LMC1 samples, 7.9% of samples were classified as positive for any STH species in Benin, 20.7% in India, and 10.8% in Malawi, while for CS1 samples, the STH positivity was 8.3%, 24.7%, and 9.0% in the same countries, respectively (**Table 1**). As this approach (DNA spiked directly into contrived sample panels) cannot account for DNA extraction inefficiency, we pursued statistical approaches to refine the qPCR Ct cut-off.

### Finite mixture models (FMM) fitted to qPCR Ct

Finite mixture models (FMM) fitted to *N. americanus* qPCR Ct estimated two Gaussian distributed sub-populations: a larger first peak (left-hand) representing the “transmissible infection” group (mean 24.1, variance 12.6) and a second peak (right-hand) representing the “indeterminate” group (mean 35.8, variance 2.8) (**Fig 2A**). Of the 5,121 samples with Ct <40 that were included in the FMM, 4,312 (84.2%) were classified as most likely (>50.0% certainty) belonging to the transmissible infection group. Within the indeterminate group, 60 and 190 samples showed >25% and >5% probability of belonging to the transmissible infection group, resulting in totals of 4,372 (85.4%) and 4,502 (87.9%), respectively. To reduce the risk of misclassification, a conservative Ct cut-off for transmissible infection was set at 34.4398, corresponding to a >5% probability of membership in the transmissible infection group. Applying this FMM cut-off, among the 2,957 samples with Ct <40 with available Kato-Katz results, 98.7% (1,495/1,514) of Kato-Katz positive samples and 79.6% (1,149/1,443) of Kato-Katz negative samples were classified as transmissible infections. When stratified by study site, the FMM-derived Ct cut-off for *N. americanus* remained consistent across all three sites. For *A. lumbricoides,* the mean and variance of the group distributions fitted by FMMs were 18.9 and 11.0 (equal variance model) for the transmissible infection population and 33.7 and 11.0 for the indeterminate population (**Fig 2B**). Among the 452 samples with a Ct <40 that were analyzed by FMM, 270 (59.6%) were classified as most likely (>50.0% certainty) belonging to the transmissible infection group. Within the indeterminate group, a further 8 and 19 samples exhibited >25% and >5% probability of membership in the transmissible infection group, resulting in totals of 278 (61.4%) and 289 (63.8%), respectively. Using the same decision criteria as used for *N. americanus*, the cut-off for >5% probability of membership in the patent infection group was defined as a Ct of 28.57587. Among the 272 samples included in the FMM derived Ct cut-off analysis for *A. lumbricoides* that also had the Kato-Katz result available, all 109 Kato-Katz positives were classified as transmissible infections. Among 163 Kato-Katz negative samples, 69 (42.3%) were classified as transmissible infections.

**Fig 2.**
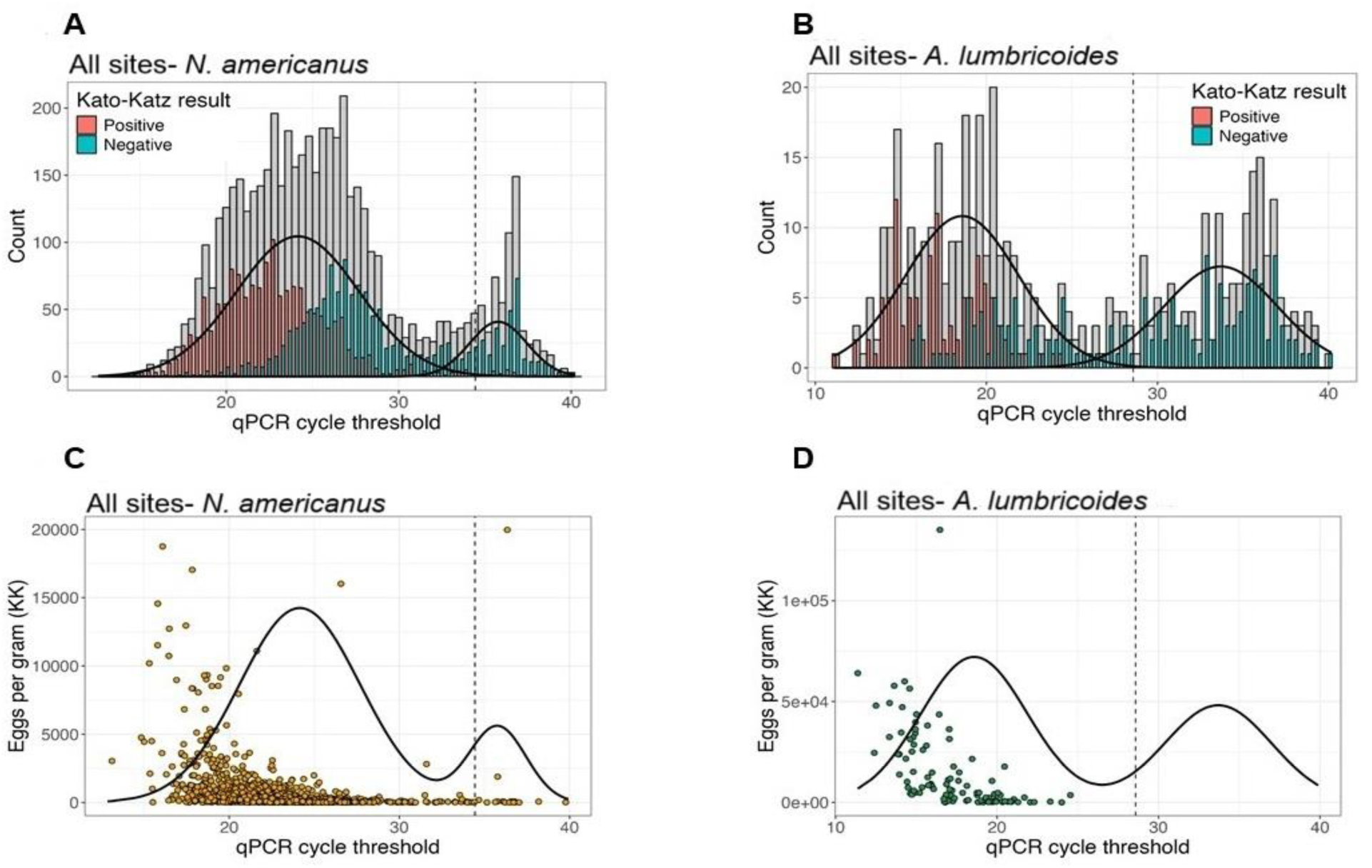
qPCR Ct values with Gaussian fit and Kato-Katz results for *Necator americanus*-and *Ascaris lumbricoides*-positive samples from DeWorm3 Trial baseline surveys conducted in Benin, India, and Malawi. The gray bar represents all samples with qPCR Ct values up to 40 for (A) *N. americanus* (n=5121) and (B) *A. lumbricoides* (n=452). Sample results are overlaid with final Gaussian distributions fitted using finite mixture models (solid black line). The dashed vertical line shows the final Ct cut-off selected to delineate samples likely to present a transmission risk. The green (Kato-Katz negative) and red (Kato-Katz positive) bars (A and B) show Ct values of all samples analyzed using Kato-Katz, disaggregated from gray bars, Panels (C) and (D) show qPCR Ct plotted against the intensity of infection as assessed by Kato-Katz.

After applying FMM-derived Ct cut-offs of 34.4398 for *N. americanus* and 28.57587 for *A. lumbricoides,* and a cut-off of 40 for *A. duodenale* and *T. trichiura*, STH positivity in LMC1 samples by qPCR was 9.7% in Benin, 24.5% in India, and 13.0% in Malawi (**Table 1**). In tested CS1 samples, STH positivity was 9.5% in Benin, 28.4% in India, and 11.7% in Malawi. After applying this FMM-derived Ct cut-offs, test positivity decreased compared to a blanket 40 Ct cut-off but was higher than with the genomic equivalent mean +3SD threshold.

### Categorization of infection intensity based on qPCR Ct distribution

For *N. americanus*, plotting infection intensity against qPCR Ct demonstrated that the majority of MHI infections, as determined by Kato-Katz, were clustered within a low Ct range, although a small minority exhibited Ct >30, including two above the final FMM derived Ct cut-off (**Fig 2C**). Similarly, for *A. lumbricoides*, infection intensity plotted against qPCR Ct showed a highly consistent pattern, where higher intensity infections assessed by Kato-Katz were strongly correlated with lower Ct (**Fig 2D**). When categorized into heavy, moderate, or light intensity infections by Kato-Katz (**Fig 3**) at each site, the median Ct for heavy-intensity infections in Benin, India, and Malawi were 17.8 (IQR 15.9 - 19.4), 18.6 (IQR 17.6 - 19.0) and 18.2 (IQR 16.6 - 18.7), respectively; for moderate-intensity the median Ct were 17.0 (IQR 15.3 - 18.3), 19.4 (IQR 18.2 - 20.4) and 18.9 (IQR 18.2 - 19.7), respectively, while samples with light-intensity infections had median Ct of 22.8 (IQR 20.9 - 24.9), 22.9 (IQR 21.0 - 25.0), and 22.4 (IQR 20.2 - 24.3), respectively. For samples that were Kato-Katz negative but positive by qPCR, median Ct values were lower - 27.4 (IQR 25.2 - 29.4), 26.9 (IQR 25.6 - 28.5), and 26.6 (IQR 24.4 - 28.4) in Benin, India, and Malawi, respectively. The data suggests that Ct distribution could be used to differentiate moderate-to heavy-intensity infections (MHI) (19.1, IQR 17.9 - 19.8) from light-intensity or Kato-Katz negative/qPCR positive infections (25.3, IQR 22.5 - 27.9) (Kruskal-Wallis test: p<0.001). For *A. lumbricoides* qPCR-positive samples in Benin, a similar trend in Ct distribution was observed from heavy-intensity (14.3, IQR 13.6 - 14.6), moderate-intensity (15.5, IQR 14.6 - 17.0), light-intensity (19.3, IQR 17.3 - 20.3) and negative by Kato-Katz/qPCR positive infections (21.4, IQR: 19.5 - 24.2) (Kruskal-Wallis test: p<0.001) (**Fig 3**). Due to low sample numbers, statistical comparisons were not made for *A. duodenale* or *T. trichiura*.

**Fig 3.**
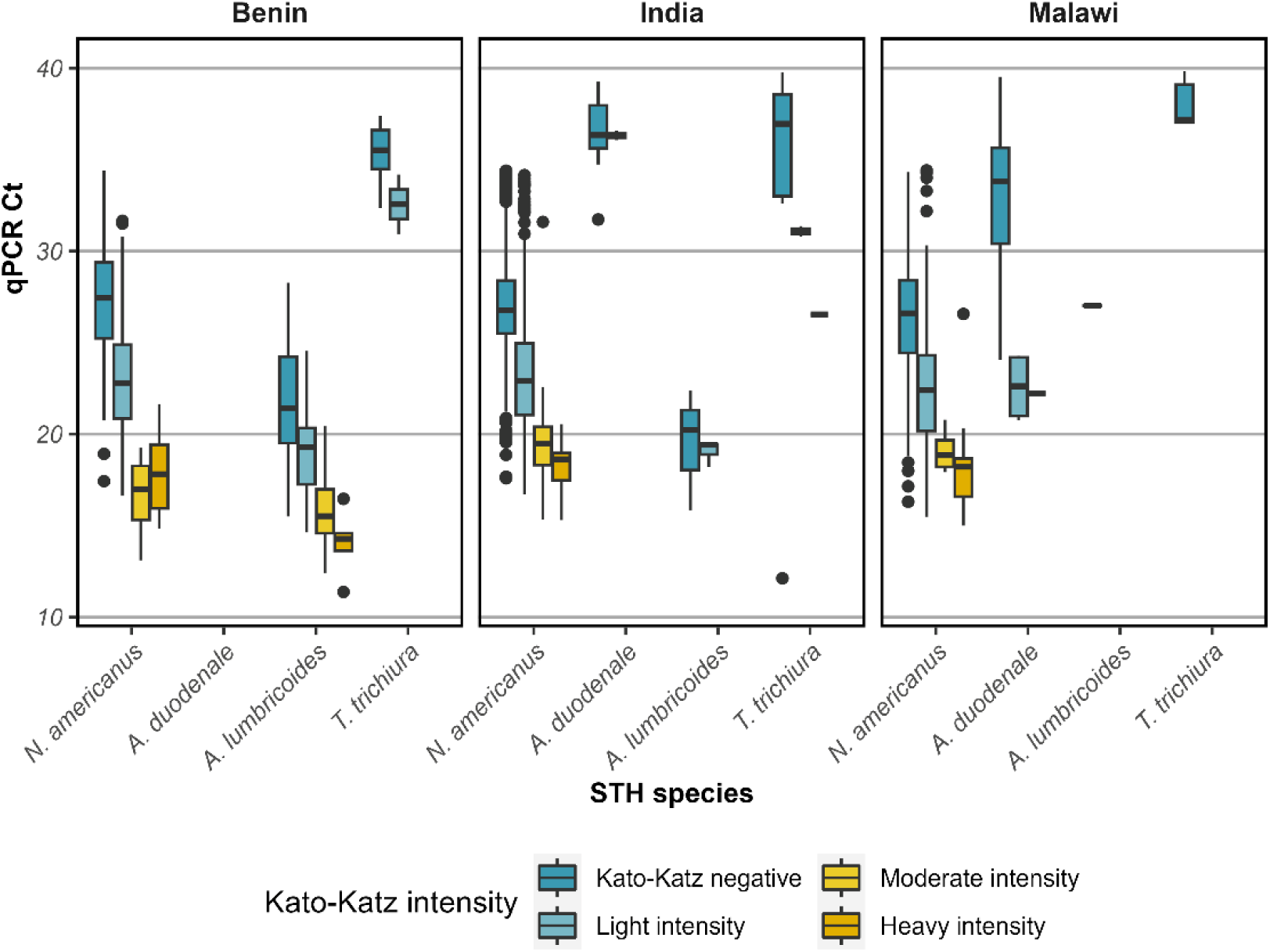
Categorization of soil-transmitted helminth Ct for Deworm3 Trial baseline survey samples from Benin, India, and Malawi stratified by intensity of infection as determined by Kato-Katz.

### Concordance and performance of Kato-Katz and qPCR tests

The concordance between double-slide Kato-Katz and qPCR with the FMM-derived Ct cut-off applied for LMC1 samples tested by both methods were 94.3%, 90.5%, and 92.1% in Benin, India, and Malawi, respectively. Also, 5.1%, 8.5%, and 6.7% of the samples from Benin, India, and Malawi that were negative for STH by Kato-Katz were positive by qPCR with an FMM-derived Ct cut-off applied (**Table 2**). Of note, a small percentage of samples (Benin: 0.6%; India: 1.0%; Malawi: 1.2%) were positive by Kato-Katz but negative by qPCR. The BLCA for the assessment of diagnostic accuracy was carried out only with samples positive for *N. americanus* and not for other species, as prevalence was low. Sensitivity, specificity, PPV, and NPV were estimated for each diagnostic test (**Table 3**). For the detection of *N. americanus* infection, the Kato-Katz technique showed a sensitivity of 73.2% (95% Bayesian Credible Interval [CrI]: 69.3 - 77.2) and a specificity of 99.4% (95% CrI: 99.2 - 99.7). In comparison, qPCR with the FMM-derived Ct cut-off showed a superior sensitivity of 96.7% (95% CrI: 94.3 - 99.0) and relatively similar specificity of 96.2% (95% CrI: 95.5 - 96.9) to the Kato-Katz.

**Table 2:**
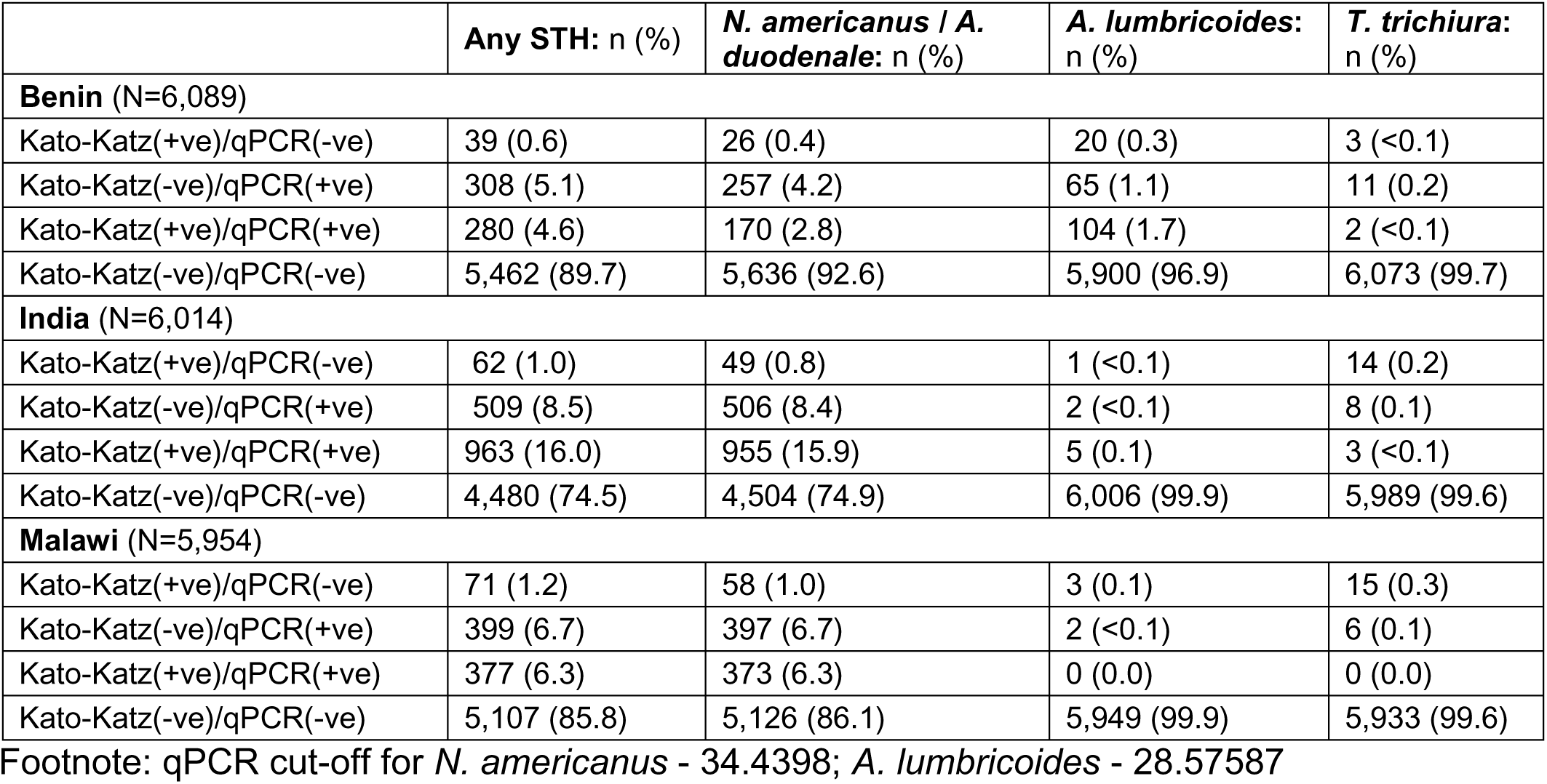
Concordance and discordance between Kato-Katz and qPCR (using FMM derived Ct cut-off) detection of soil-transmitted helminth infections.

**Table 3:** Sensitivity and specificity of Kato-Katz and qPCR for the detection of Hookworm* using Bayesian latent class analysis.

| Diagnostic methods | Kato-Katz<br>% (95% CrI) | qPCR<br>% (95% CrI) |
| --- | --- | --- |
| Sensitivity | 73.2% (69.3% - 77.2%) | 96.7% (94.3% - 99.0%) |
| Specificity | 99.4% (99.2% - 99.7%) | 96.2% (95.5% - 96.9%) |
| PPV | 94.4% (92.0% - 96.9%) | 77.1% (73.0% - 81.1%) |
| NPV | 96.6% (95.8% - 97.2%) | 99.5% (99.2% - 99.9%) |

Footnote: \*Hookworm includes only *N. americanus* qPCR data. Benin n=6,089; India n=6,014; Malawi n=5,954; PPV – Positive Predictive Value; NPV – Negative Predictive Value; CrI – Bayesian Credible Interval

## Discussion

As large-scale operational research efforts, such as the DeWorm3 cluster randomized trial and other studies [32, 33] explore the potential for STH transmission interruption and/or achieving morbidity elimination [34, 35], there is a growing need to interpret available qPCR diagnostic test results to assess public health outcomes and impact of interventions. Numerous publications compare multiple coproscopic techniques [9, 36–39] or evaluate various microscopy-based techniques against PCR or other molecular detection methods [11, 12, 40, 41], but with very few exceptions [42, 43], the published literature is largely void of studies comparing various qPCR assays. Different qPCR assays are no more similar in performance than are different microscopy-based techniques, with variations in efficiency, linear dynamic range, analytical and clinical limits of detection, and precision [44]. Accordingly, appropriate Ct cut-offs must be experimentally determined for each assay used rather than using a blanket Ct <40 [45]. A combination of wet lab experimentation and statistics were used to investigate the relationship between qPCR Ct distribution and detection of transmissible infections. Due to extraction efficiency concerns with high variability observed in the single-egg spiking experiments and an inability to account for extraction using genome-equivalent spiking experiments, a statistical approach was used to determine Ct cut-off.

While multiple studies employing different qPCR assays with unique primer target sequences have shown that, for STH detection, qPCR Ct values show a bimodal distribution [12, 18], to our knowledge, this study is the first to address this bimodal distribution, explore its significance for interpretation, and apply it in the context of a clinical trial [35]. We hypothesized that the first peak on the Ct density plot corresponded to transmissible egg positive infections, while the second peak corresponded to detection of nucleic acid from either degraded ova or shed helminth material, or from unpaired (single sex) helminth infections, none of which contribute to transmission. Based on this interpretation, the implementation of Ct cut-offs was proposed to differentiate between transmissible and non-transmissible infections. The FMM statistical modeling approach has been applied to serological response data from populations with unknown degrees of exposure to antigens of interest [46–48] and, less commonly, for determining Ct cut-offs for qPCR-based assays [49]. In the context of the DeWorm3 Trial, FMM-derived Ct cut-off values captured over 98% of samples that were Kato-Katz positive for *N. americanus* and 100% of samples that were Kato-Katz positive for *A. lumbricoides*. Additionally, a significant number of samples that were classified as negative by Kato-Katz were identified as qPCR positive using this approach. In our analysis, the FMM derived Ct cut-offs stratified by three sample collection sites and by two primary testing laboratories were found to be similar, despite differences in prevalence and intensity distribution. This indicates that when robust cut-offs are established for a given assay, they could be applied in multiple settings or studies.

Accurately determining the sensitivity and specificity of STH diagnostic methods, both coproscopic and molecular, remains challenging due to the absence of a true gold standard [50]. In this study, we used BLCA and found qPCR to be more sensitive than double-slide Kato-Katz, which is consistent with previous studies [8, 11]. It is also important to consider that when implemented during monitoring and evaluation of programs in a “real-world” setting, Kato-Katz has greater variability and reduced sensitivity compared to a controlled “study” setting [51]. While both techniques require skilled personnel to perform, unlike microscopy-based methods, where results require interpretation by skilled personnel, qPCR result does not rely on human interpretation. Accordingly, the true difference in sensitivity between methods may be even greater than the reported difference. Despite the improved sensitivity of detection, considerable focus remains on providing programmatically relevant information and effectively correlating qPCR Ct with infection intensity categories defined by EPG [40, 52]. In this study, we found that MHI for *N*. *americanus* and *Ascaris* could be delineated from light-intensity infections or negative Kato-Katz results. This finding aligns with results observed in a limited number of other studies [12, 41] and suggests that Ct value can be used to assess infection intensity, at least as a means of “binning” infections into programmatically relevant categories. Given these findings, a full understanding of performance characteristics is warranted.

A small percentage of the samples (≤1%) were positive by Kato-Katz and negative by qPCR. This discordance was minimal and to be expected in large-scale studies and occurred across all three study sites and for all STH species [53, 54]. Kato-Katz positive, qPCR negative samples may represent Kato-Katz false-positives, as evidence suggests that microscopy-based techniques are prone to specificity challenges when compared with molecular methods [12, 55–57]. In this study setting, the discordance was also a result of extraction inefficiency in a small number of samples, due to the automated extraction protocol used, and is the limitation of this high throughput protocol [22]. The Ct values detected from fecal sample DNA extracts can be influenced by multiple factors [58, 44]. Degraded or low DNA content, heterogeneity in egg distribution, inefficient lysis of ova, loss in sensitivity due to multiplexing, or the presence of inhibitors may also have impacted the results [10, 12, 59–64]. Another limitation of this study is that qPCR, although performed at the other sites, was not feasible in Malawi due to challenges in accessing advanced laboratory infrastructure and maintaining a reliable reagent supply chain.

In conclusion, while qPCR has proven to be highly sensitive in detecting STH infections, especially in light-intensity settings, with the development of extremely sensitive assays targeting repetitive sequences, certain research questions may render it important to differentiate epidemiologically relevant transmissible infections from indeterminate samples, which are unlikely to represent a transmissible infection. Accordingly, we have developed an evidence-based approach, including assay characteristics, experimental investigation, and statistical models, to determine qPCR Ct cut-offs.

## Supporting information

**S1 Table: Egg spiking experimental approaches to determine qPCR Ct cut-offs for soil-transmitted helminth infections**

**S2 Table: Genome-equivalent spiking experiment-derived qPCR Ct cut-off**

**S1 Fig:**
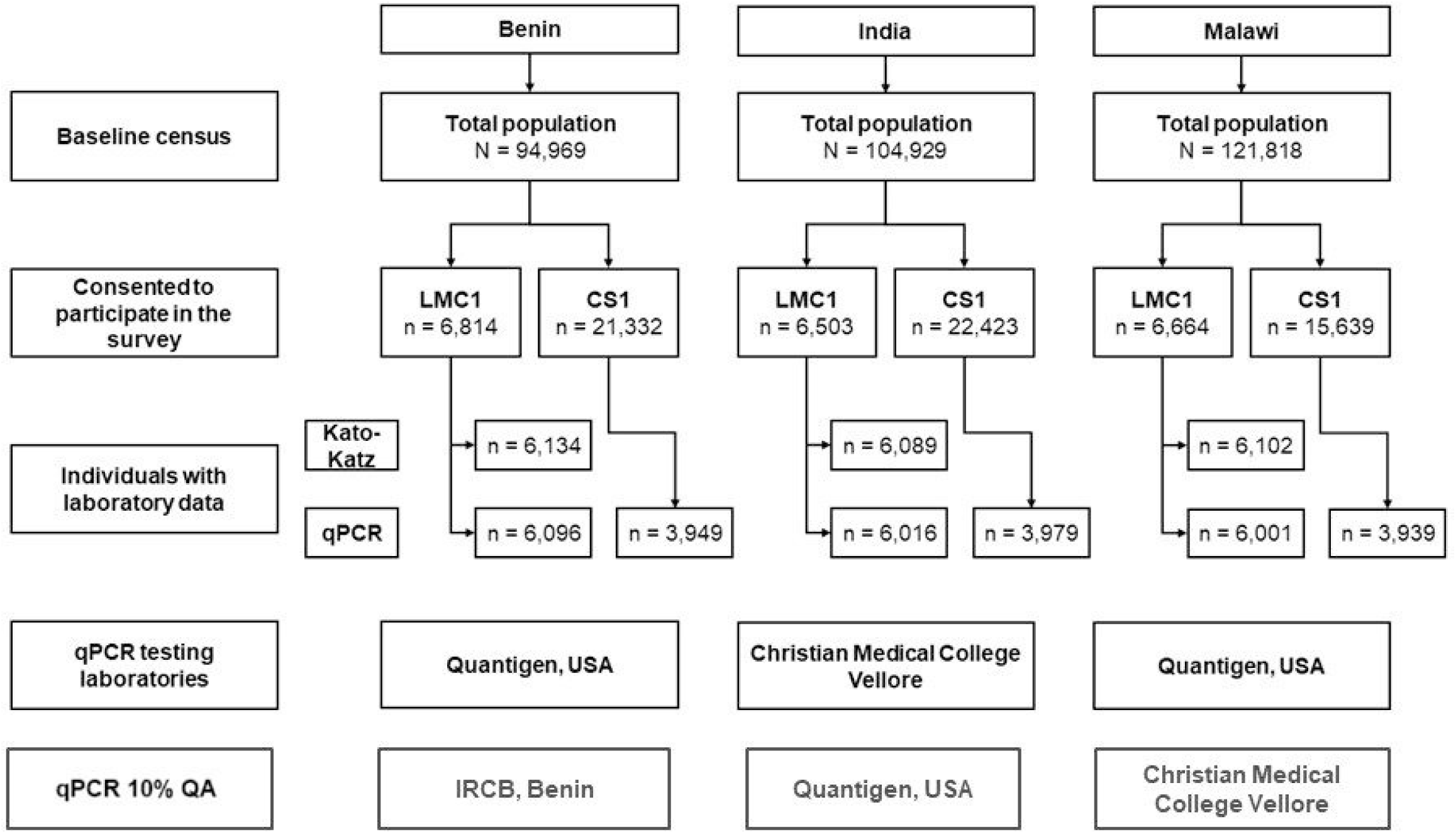
Flow chart of the DeWorm3 Trial baseline surveys sample collection and testing sites for Kato-Katz and qPCR.

## Acknowledgements

The authors wish to thank all of the study participants and communities, ministries of health and other local, regional, and national partners who have participated or supported the DeWorm3 study. We would specifically like to acknowledge Jaco J Verweij, Lisette van Lieshout, Brandon Troy Leader, and Leah Padgett for their guidance during the validation of the qPCR assay. We would like to acknowledge the work of all members of the DeWorm3 study teams and affiliated institutions, in particular the laboratory staff involved in qPCR testing at Quantigen, Fishers, IN, Smith College, Northampton, MA, Christian Medical College, Vellore, India, and the Institut de Recherche Clinique du, Benin. The data from the India site is part of the PhD research work undertaken by Malathi Manuel under the guidance of Sitara Swarna Rao Ajjampur, registered with The Tamil Nadu Dr. M.G.R. Medical University, Chennai. We also wish to acknowledge the contributions of the DeWorm3 Data Safety Monitoring Committee, including Richard Hayes, Harriet Mpairwe, Purushothaman Jambulingam, and Jimmy Whitworth.

## Data availability statement

Data are available from the DeWorm3 Institutional Data Access Committee (contact via, https://depts.washington.edu/deworm3/) and in Vivli, the data sharing platform (https://doi.org/10.25934/PR00010754) for researchers who meet the criteria for access to these data.

## Author contributions

**Conceptualization:** Nils Pilotte, Rachel Pullan, Steven Williams, Judd L Walson, Sitara SR Ajjampur

**Data curation:** Sean Galagan

**Formal analysis:** Joseph W.S. Timothy, Rachel Pullan, Sean Galagan, Gideon John Israel

**Funding acquisition:** Judd L Walson

**Investigation:** Nils Pilotte, Malathi Manuel, Craig T Connors, Victor Omballa, Monica Pechanec Voss, Ushashi C Dadwal, Andrew M Gonzalez, Justine Ahlonsou, David Chaima, Zayina Zondervenni Manoharan

**Methodology:** Nils Pilotte, Monica Pechanec Voss, Doug Rains, Joseph W.S. Timothy, Malathi Manuel, Sitara SR Ajjampur

**Project administration:** Nils Pilotte, Malathi Manuel, Monica Pechanec Voss, Doug Rains, Steven A. Williams, Judd L Walson, Sitara SR Ajjampur

**Resources:** Doug Rains, Steven A. Williams, Rachel Pullan, Judd L Walson, Sitara SR Ajjampur

**Supervision:** Doug Rains, Kristjana H. Ásbjörnsdóttir, Steven A. Williams, Khumbo Kalua, Adrian J F Luty, Moudachirou Ibikounlé, Robin Bailey, Rachel Pullan, Judd L Walson, Sitara SR Ajjampur

**Visualization:** Joseph W.S. Timothy, Sean Galagan, Gideon John Israel, Rachel Pullan

**Writing** – original draft: Malathi Manuel, Nils Pilotte, Sitara SR Ajjampur

**Writing – review & editing**: Malathi Manuel, Nils Pilotte, Joseph W.S. Timothy, Sean Galagan, Gideon John Israel, Craig T Connors, Victor Omballa, Monica Pechanec Voss, Ushashi Dadwal, Andrew M Gonzalez, Justine Ahlonsou, David, Zayina Zondervenni Manoharan, Doug Rains, Kristjana H. Ásbjörnsdóttir, Steven A. Williams, Khumbo Kalua, Adrian J F Luty, Moudachirou Ibikounlé, Robin Bailey, Rachel Pullan, Judd L Walson, Sitara SR Ajjampur

